# Systemic Effects of Missense Mutations on SARS-CoV-2 Spike Glycoprotein Stability and Receptor Binding Affinity

**DOI:** 10.1101/2020.05.21.109835

**Authors:** Shaolei Teng, Adebiyi Sobitan, Raina Rhoades, Dongxiao Liu, Qiyi Tang

**Author notes:** Corresponding Authors. Address correspondence to: Shaolei Teng and Qiyi Tang.

## Abstract

The spike (S) glycoprotein of SARS-CoV-2 is responsible for the binding to the permissive cells. The receptor-binding domain (RBD) of SARS-CoV-2 S protein directly interacts with the human angiotensin-converting enzyme 2 (ACE2) on the host cell membrane. In this study, we used computational saturation mutagenesis approaches, including structure-based energy calculations and sequence-based pathogenicity predictions, to quantify the systemic effects of missense mutations on SARS-CoV-2 S protein structure and function. A total of 18,354 mutations in S protein were analyzed and we discovered that most of these mutations could destabilize the entire S protein and its RBD. Specifically, residues G431 and S514 in SARS-CoV-2 RBD are important for S protein stability. We analyzed 384 experimentally verified S missense variations and revealed that the dominant pandemic form, D614G, can stabilize the entire S protein. Moreover, many mutations in N-linked glycosylation sites can increase the stability of the S protein. In addition, we investigated 3,705 mutations in SARS-CoV-2 RBD and 11,324 mutations in human ACE2 and found that SARS-CoV-2 neighbor residues G496 and F497 and ACE2 residues D355 and Y41 are critical for the RBD-ACE2 interaction. The findings comprehensively provide potential target sites in the development of drugs and vaccines against COVID-19.

## INTRODUCTION

Coronaviruses are a large family of enveloped RNA viruses typically transmitted from animals to animals. These viruses are usually found in bats, birds, and mammals (1), but seven coronavirus members are found to infect humans. The paradigm shift in the mode of transmission of three pathogenic coronaviruses, including severe acute respiratory syndrome coronavirus 2 (SARS-CoV-2), severe acute respiratory syndrome (SARS-CoV) and Middle East respiratory syndrome coronavirus (MERS-CoV), has resulted in recent outbreaks. The SARS-CoV epidemic began in Shunde, Guangdong, China, in November 2002. It ended in 2003 with more than 8000 cases in over 28 countries and ~800 deaths. The presence of a SARS-CoV-like coronavirus in bats indicated that the natural reservoir of SARS-CoV is horseshoe bat (2, 3). The MERS-CoV epidemic began in Saudi-Arabia in June 2012, and the transmission of MERS-CoV crossed the Arab peninsula to nearby countries. Scientists revealed that the Arabian camel (dromedaries) is the reservoir host to MERS-CoV, and they also spread the virus to humans. As of January 2020, the World Health Organization (WHO) reported 2,519 MERS-CoV cases in 27 countries and ~866 deaths. Although both SARS-CoV and MERS-CoV were transmitted from animals to humans, human to human transmission was found plausible (2). The most recent outbreak of Coronavirus Disease 2019 (COVID-19) is caused by a novel coronavirus, SARS-CoV-2. The virus is closely related to the SARS-like CoV, RaTG13 virus which is found in bats (4). SARS-CoV-2 is so far the most pathogenic coronavirus. As of May 23, 2020, WHO reported over 5 million confirmed cases of COVID-19 in 216 countries, resulting in > 331,000 deaths.

The spike (S) glycoprotein is critical in terms of the virulence of pathogenic coronaviruses. The homotrimeric protein is responsible for mediating virus entry of SARS-CoV-2 via the Angiotensin-converting enzyme 2 (ACE2) receptor on the host cell membrane (5, 6). Thus, the analysis of SARS-CoV-2 S protein is a high research priority for vaccine design against COVID-19. The S protein consists of an S1 and an S2 subunit. Proteolytic cleavage at the boundary of the S1 and S2 subunits is activated by the binding of the S1 subunit with the ACE2 receptor (5). The pair-wise alignment of SARS-CoV2 and SARS-CoV spike protein revealed a ~55% identity in the S1 subunits and ~91% identity in the S2 subunit (7). This subsequent disassociation allows for a conformation change in the S2 subunit that allows it to fuse with the SARS-CoV-infected cell membrane (8). The receptor-binding domain (RBD) of S1 includes a core and a receptor-binding motif (RBM) that specifically recognizes ACE2. The amino acid sequence similarities of SARS-CoV and SARS-CoV-2 in the S protein, RBD, and RBM are 76%, 73% and 50%, respectively(9).

The interactions between RBD and ACE2 are critical for the host range and cross-species infections of SARS-CoV. A simulation study showed that SARS-Cov-2 S protein has a higher affinity to ACE2 compared to that of Bat-CoV S to the ACE2 receptor (10). An investigation of the RBD of SARS-CoV-2 demonstrated that SARS-CoV-2 RBD had a significantly greater binding affinity to the human ACE2 (hACE2) and bat ACE2 receptors (11). A recent study showed that the binding affinity of SARS-CoV-2 to ACE2 is comparable to that of SARS-CoV, as plasma containing antibodies to the SARS-CoV reduced the transduction efficiency of SARS-CoV-2 in cells (5).

Structure analyses revealed the atomic details of SARS-Cov and SARS-Cov-2 S proteins and the binding interface between RBD and ACE2. The conformational state of the C-terminal domain 1 (CTD1) of SARS-CoV S1 is responsible for the binding of S to the human ACE2 (hACE2) receptor. SARS-CoV S has an inactive state with a symmetric conformational state representing the CTD1 in the “down” position and an active state with three asymmetric conformational states representing CTD1 in the “up” position (12). RBD is also the key determinant of cross-species and human-to-human transmissibility of SARS-CoV (13). The SARS-CoV S1 subunit is divided into the N-terminal domain and the receptor-binding domain (RBD). The RBD attaches to the peptidase domain of the ACE2 receptor via its RBM. Recent studies revealed the similar modes of SARS-Cov-2 S conformational states and RBD-ACE2 binding. A prefusion ectodomain trimer in the opened and closed conformational states of the SARS-COV-2 S was determined using the cryogenic-EM study (5). In the open conformational state, SARS-CoV-2 S protein can recognize the ACE2 to initiate the viral entry. Crystallographic study showed that SARS-CoV-2 spike RBD can bind hACE2 via its RBM (14). Want et al. were able to demonstrate through immunostaining that the CTD of the SARS-CoV-2 S1 co-localizes with the hACE2, demonstrating that this region contains the RBD that mediates the attachment of the virus to the host cell membrane (14). They demonstrated that 15 amino acids of the RBD of SARS-CoV-2 S1 established more atomic bonds with the hACE2 when compared to the crystallized structures of the RBD of SARS-CoV S1 in complex with the ACE2 receptor. These interface variations are thought to alter the affinity of the RBD for the ACE2 receptor. The ACE2 receptor consists of an N-terminal peptidase domain, a C-terminal Collectrin-like domain, a transmembrane helix, and an intracellular domain. The crystal structures of the ACE2 receptor in complex with the amino acid transporter B0AT1 showed that ACE2 forms homodimers mediated primarily by a ferredoxin-like fold domain, located between the peptidase domain and the transmembrane helix (15).

The genome of RNA virus can easily generate mutations as virus spreads. Genetic studies showed that SARS-CoV has 18-fold increases in mutation rates compared to other RNA viruses (16). SARS-CoV-2 spreads rapidly worldwide and accumulates many mutations. The constant emergence of new mutations in SARS-CoV-2 is the major challenge for the ongoing development of antiviral drug and broad neutralizing antibodies. The *in silico* methods can readily quantify the effects of coding mutations on protein function and structure. A bioinformatics analysis based on the homology modelling of SARS-CoV-2 predicted that the mutations in the RBD region reduce the binding energy between the RBD and the ACE2 (10). However, a comprehensive effect map of SARS-CoV-2 S mutations still lacks for identifying the target sites for vaccine design. In-silico saturation mutagenesis provides a fast methodology to investigate all possible mutations and identify the potential functional sites. It has been applied to Mycobacterium research to determine the effects of missense mutations on thermostability, affinity, conformational changes and protein-protein interaction (17). The infection of SARS-CoV-2 in permissive cells is largely depending on the interaction of the viral S protein and ACE2. The affinity of the S and ACE2 and the stability of RBD-ACE2 complex are hence important to be investigated. We have applied the structure-based tools to quantitatively assess the effects of damaging mutations on protein stability and protein-protein interaction (18–20). In the present study, we applied saturation mutagenesis to investigate 18,354 missense mutations in SARS-CoV-2 S, 3,705 mutations in SARS-CoV-2 RBD, and 11,324 mutations in hACE2. We applied structure-based energy calculations to quantify the systemic effects of missense mutations on the stability of and binding affinity of the S and hACE2 proteins. We studied the mutations involved in N-linked glycosylation which is important for the stability of SARS-CoV-2 S protein (21). We identified the potential target sites critical for the designs of antiviral drugs and vaccines against COVID-19.

## MATERIALS AND METHODS

### Structure preparation

All structures are collected from Protein Data Bank (PDB) (22). We collected the structure of the SARS-CoV-2 2P (P980-P987) spike (S) ectodomain glycoprotein in the open state (PDB ID: 6VYB). To study the protein stability of wide type SARS-CoV-2 S, we separated the A chain of 6VYB and introduced P986K and P987V mutations using FoldX (23). The S receptor-binding domain (RBD) complexed with its receptor ACE2 (PDB ID: 6LZG) was used to investigate the RBD-ACE2 binding affinity and protein stability of SARS-CoV-2 RBD and hACE2. The SARS-CoV-2 S structure in closed state (6vxx), SARS-CoV S glycoprotein (6acg) and RBD-ACE2 complex (2ajf), and MERS-COV S protein (5w9j) were collected for the comparison studies. The PyMOL (http://www.pymol.org/) was used to generate protein structural images and perform structure alignments.

### Mutation collection

To determine the effects of the mutations on protein structure, we collected all mutations that can be mapped onto the corresponding structures. Virus variations of SARS-CoV-2 were collected from 2019 Novel Coronavirus Resource (2019nCoVR) from China National Center for Bioinformation (https://bigd.big.ac.cn/ncov). As of 04/30/2020, we collected 385 viral missense variations in 325 residue positions of SARS-CoV-2 S protein.

### Sequence Analysis and Mutation Pathogenicity

The amino acid sequences of S protein of SARS-CoV-2 (Entry: P0DTC2), SARS-CoV (Entry: P59594) and MERS-CoV (Entry: K9N5Q8) and human ACE2 (Entry: Q9BYF1) were downloaded from UniProt (24). The pairwise sequence alignment was carried out using EMBOSS Water multiple sequence alignment was performed using Clustal Omega and (25).

The protein sequences were submitted to SNAP (26) to predicts the effect of mutations on protein function for SARS-CoV-2 S and hACE2. SNAP uses neural networks to integrate many sequencebased bioinformatics tools to analyze the mutation pathogenicity. It takes a protein sequence as input and gives a reliability index (RI) as a measure of the prediction accuracy. The prediction scores range from −100 (neutral) to 100 (effect). Positive scores indicate that the mutations have damaging effects on protein function.

### Free Energy Changes Calculations

The effects of mutations on protein stability of S proteins and binding affinity of RBD with hACE2 were estimated by the change of the folding free energy (ΔΔG) and the change of the binding free energy (ΔΔΔG) between the mutant structure (MUT) and wild-type (WT) structure, respectively. FoldX (23) was used for energy calculations. It can be used to perform mutagenesis and predict protein stability and biding changes as well as calculate energy contributions of total energy, electrostatic, Van der Waals interactions and other energy terms. Before performing any Foldx analysis, all protein structures were repaired using the ‘RepairPdb’ command, which works by mutating certain residues to themselves to reduce the overall free energy of the protein structure.

For stability analysis, the ‘BuildModel’ command was used. This command takes as input, both repaired protein structure and a list of point mutations. The output computes energy changes brought about by each point mutation. The advantage of the reference wildtype model is that it considers the neighboring side chains that are moved in the process of generating the mutant protein. The change of the folding free energy (ΔΔG) introduced by an amino acid substitution is an indicator of how this mutation affects the stability of the structure. It was calculated using:

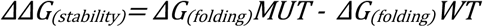

A negative *ΔΔG* value suggests that the mutation can stabilize the monomer protein and a positive value of *ΔΔG* indicates that it makes the protein unstable.

For interaction analysis, the ‘AnalyseComplex’ command was carried out, where the user can calculate the binding affinity of a protein structure. This feature determines the interaction of proteins by unfolding each protein and evaluating their individual energies. The program then subtracts the individual protein energies from the energy of the bound proteins or protein complex. The change of binding energy (ΔΔΔG) between the mutant and wild-type structures was used to estimate the effect of the missense mutations on the binding affinity of RBD-ACE2 protein complex. It was computed by:

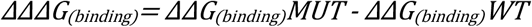

A negative ΔΔΔ*G* value suggests that the mutation strengthens the binding affinity, whereas a positive value of ΔΔΔ*G* indicates that the mutation weakens the protein-protein interaction.

R package (https://www.r-project.org/) was used to generate graphs and perform ANOVA test and t test in statistical comparisons of ΔΔG and ΔΔΔG values and SNAP scores.

## RESULTS

### Effects of mutations on SARS-CoV-2 full-length S protein stability (ΔΔG)

Since S protein is critical for SARS-CoV-2 infection, we want to anatomize its amino acids that affect its stability. We used the cryo-EM structure of the SARS-CoV-2 spike ectodomain structure in the open state (PDB ID: 6vyb) as the SARS-CoV-2 full-length S structure. This structure covers 27-1141 amino acid region and includes 966 residues (76%) of SARS-CoV-2 full-length S protein. To determine the effects of the systematic mutations on SARS-CoV-2 S stability, we generated 18,354 mutations by mutating all 966 residues to all other 19 amino acid types and computed the changes of the folding free energy (ΔΔG) introduced by these mutations. Of 18,354 mutations, 11,313 (61.6%) mutations can destabilize SARS-CoV-2 full-length S protein (ΔΔG > 0.5 kcal/mol), 1,942 (10.6%) can stabilize the protein (ΔΔG < −0.5 kcal/mol) and the remaining 5,099 (27.8%) have no effects on SARS-CoV-2 S stability (0.5 > ΔΔG > −0.5 kcal/mol). As shown in Figure 1a, 31.3% mutations have strong effects (ΔΔG > 2.5 kcal/mol) and 30.4% mutations have moderate effects (0.5 < ΔΔG <= 2.5 kcal/mol) on protein destabilization. The heatmap of in Figure 2A also suggests that the most mutations can decrease SARS-CoV-2 full-length S protein stability. In contrast, only 0.2% mutations have strong effects (ΔΔG < −2.5 kcal/mol) and 10.4% mutations have moderate effects (−2.5 <= ΔΔG < −0.5 kcal/mol) on S protein stabilizing (Figure 1A). We also calculated the folding energy changes based on the cryo-EM structure of the SARS-CoV-2 S in the closed state (PDB ID: 6vxx) and found that the results are highly correlated with those in the open state. As shown in Supplementary Figure 2, Pearson’s correlation coefficient (R) values of ΔΔG for full-length S in the open and closed conformations are 0.9192 and 0.9509 for all mutations and residue mean values, respectively.

**Figure 1.**
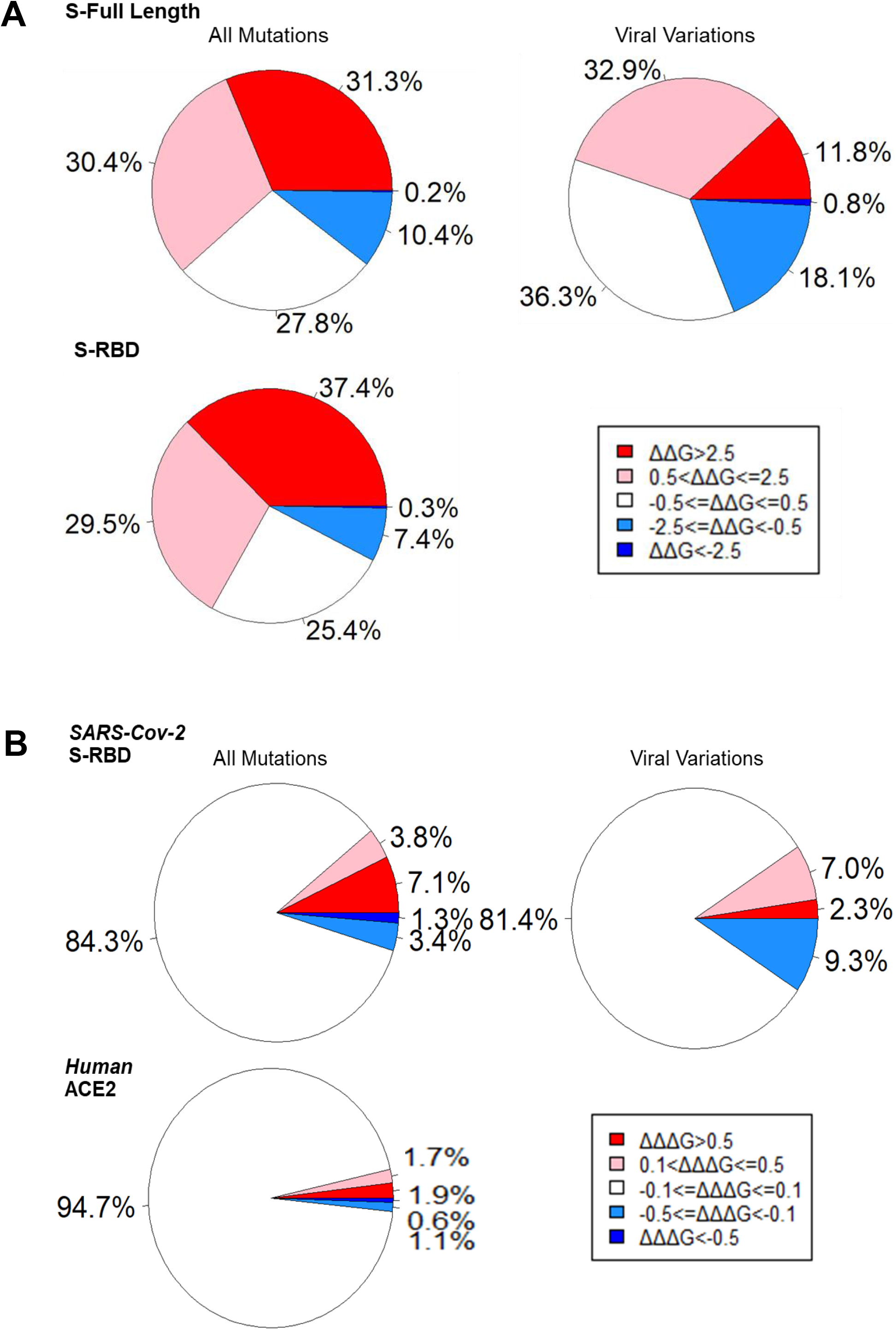
Pie charts summarize the contribution of (**A**) the stability effects of all mutations and viral variations on SARS-Cov-2 Full-Length S and all mutations on RBD and (**B**) the binding affinity effects of SARS-Cov-2 RBD all mutations and viral variations and all Human ACE2mutations.

**Figure 2.**
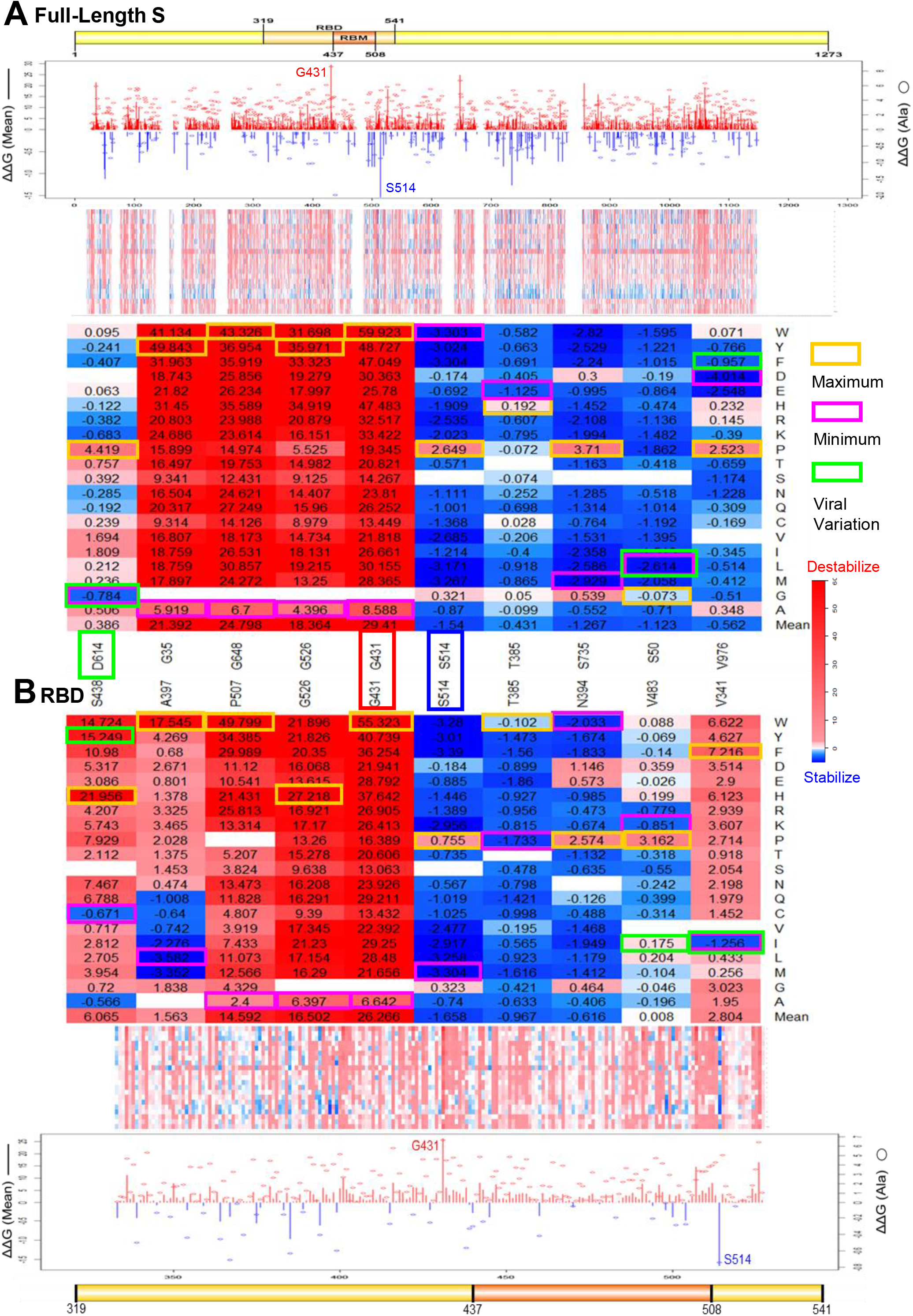
Effects of residues and mutations in SARS-Cov-2 (**A**) full-length S and (**B**) RBD on protein stability. Line charts summarize the folding energy changes for ΔΔG mean of residues (bar) and ΔΔG of substitutions to Alanine (circle). Heatmaps show the ΔΔG of all mutations and the mutations in key residues. Maximum (yellow) and Minimum (magenta) ΔΔG values are labelled for each residue position. The ΔΔG values of viral variations are shown in green boxes. The key residues G431 and K514 are marked in the line charts and heatmaps.

We present the key residues and mutations with significant effects on SARS-CoV-2 S protein stability in Table 1 and Figure 2. As the line chart shown in Figure 2A, the mean value of folding change (ΔΔG) in protein stability at each SARS-CoV-2 full-length S residue position ranged from 29.41 kcal/mol in G431 to −1.54 kcal/mol in S514. We listed the amino acid positions that strongly stabilize or destabilize spike protein in the Table 1 according to the mean values of ΔΔG of mutations in each position. The mutations at glycine residues G431, G648 and G35 residues have maximum destabilizing effects on the full-length S protein. In contrast, the mutations at serine residues S514, S735 and S50 residues have induced highly stabilizing effects on the overall protein. Noteworthily, both G431 with maximum ΔΔG and S514 with minimum ΔΔG are located in the RBD region. Moreover, residues G526 (ΔΔG mean = 18.364 kcal/mol) and T385 (ΔΔG mean = −0.431 kcal/mol) in RBD have significant effects on the stability of full-length S protein. The mutations with maximum and minimum ΔΔG values are listed in Table 1. Among all mutations destabilizing full-length S protein, the glycine mutation G431W has introduced the largest positive folding energy at ΔΔG = 59.92 kcal/mol. However, V976D has the minimum ΔΔG at −4.014 kcal/mol and most of the mutations in V976 (ΔΔG mean = −0.562 kcal/mol) can increase the stability of full-length S protein (Figure 2a).

**Table 1.** Effects of key residues and mutations on SARS-Cov-2 S stability.

| Residue | $\Delta\Delta G$ (Mean) | Mutation with Maximum $\Delta\Delta G$ | | Mutation with Minimum $\Delta\Delta G$ | |
| --- | --- | --- | --- | --- | --- |
| *Full-Length S* |  |  |  |  |  |
| **G431** | **29.41** | **G431W** | **59.923** | G431A | 8.588 |
| G648 | 24.798 | G648W | 43.326 | G648A | 6.700 |
| G35 | 21.392 | G35Y | 49.843 | G35A | 5.919 |
| G526 | 18.364 | G526Y | 35.971 | G526A | 4.396 |
| D614 | 0.386 | D614P | 4.419 | <u>D614G</u> | <u>-0.784</u> |
| **S514** | **-1.54** | S514P | 2.649 | S514F | -3.304 |
| S735 | -1.267 | S735P | 3.710 | S735M | -2.929 |
| S50 | -1.123 | S50G | -0.073 | S50L | -2.614 |
| V976 | -0.562 | V976P | 2.523 | **V976D** | **-4.014** |
| T385 | -0.431 | T385H | 0.192 | T385E | -1.125 |
| *RBD* |  |  |  |  |  |
| **G431** | **26.266** | **G431W** | **55.323** | G431A | 6.642 |
| G526 | 16.502 | G526H | 27.218 | G526C | 9.39 |
| P507 | 14.592 | P507W | 49.799 | P507A | 2.400 |
| S438 | 6.065 | S438H | 21.956 | S438C | -0.671 |
| V483 | 0.008 | V483P | 3.162 | V483R | -0.779 |
| **S514** | **-1.658** | S514P | 0.755 | S514F | -3.390 |
| T385 | -0.967 | T385W | -0.102 | T385P | -1.733 |
| N394 | -0.616 | N394P | 2.574 | N394W | -2.033 |
| A397 | 1.563 | A393W | 17.545 | **A397L** | **-3.582** |
| V341 | 2.804 | V341F | 7.216 | V341I | -1.256 |

The residues with the largest or smallest $\Delta\Delta G$ means and mutations with the largest or smallest $\Delta\Delta G$ values in all predictions are shown as bold fonts. The dominant pandemic form D614G is underlined.

### SARS-CoV-2 RBD mutations important for domain stability

The full-length S protein is cleaved to S1 and S2 by cellular protease, furin. S1 binds to ACE2 through its RBD. Hence, we wonder whether S1 RBD stability could be affected by amino acid substitutions. In doing so, we used the crystal structure of SARS-CoV-2 S RBD complexed with its receptor ACE2 (PDB ID: 6lzg) and analyzed a total of 3,705 mutations generated from 195 residues of its RBD chain. Of 3,705 mutations, 2,478 (66.9%) mutations can destabilize SARS-CoV-2 RBD protein (ΔΔG > 0.5 kcal/mol), 285 (7.7%) can stabilize the protein (ΔΔG < −0.5 kcal/mol), and the remaining 942 (25.4%) have no effects on SARS-CoV-2 S stability (−0.5 <= ΔΔG <= 0.5 kcal/mol). Consistent to the analysis results of the full-length S protein, there are more mutations causing destabilization than stabilization of RBD (Figure 1A). 37.4% of RBD mutations have strong effects (ΔΔG > 2.5 kcal/mol) on protein destabilizing compared to 31.3% mutations with strong effects on full-length S protein. Interestingly, there are fewer RBD mutations (7.4%) that have moderate effects (−2.5 <= ΔΔG < −0.5 kcal/mol) on protein stabilizing, compared to 10.4% those mutations in full-length S protein. In addition, we showed that the effects of mutations on RBD stability are highly correlated with those on full-length S protein (Supplementary Figure 1B). The Pearson’s correlation coefficient (R) values of ΔΔG on RBD and full-length S are 0.8322 and 0.8733 for all mutations and residue mean values, respectively. This finding suggests that the energy calculations based on the crystal structure of RBD-ACE2 complex (PDB ID: 6lzg) are consistent with those based on the cryo-EM structure of full-length S structure (PDB ID: 6vyb).

The top RBD residues and mutations with maximum and minimum ΔΔG values are shown in Table 1 and Figure 2B. The average change in protein stability at each SARS-CoV-2 S residue position ranged from 26.266 kcal/mol in G431 to −1.658 kcal/mol in S514. Notably, G431 and S514 are also the top residues altering the full-length S stability. Interestingly, G431A has the maximum ΔΔG values among all substitutions to alanine in both full-length S and RBD, and S514A has the minimum ΔΔG value among all substitutions to alanine in RBD. These results indicate that the side-chains of G431 and S514 have important contributions to the RBD stability (Figure 2). Other residues strongly destabilizing RBD include G526 (ΔΔG mean = 16.502 kcal/mol) and P507 (ΔΔG mean = 14.592 kcal/mol). In contrast, the mutations at residue positions T385 (ΔΔG mean = −0.967 kcal/mol) and N394 (ΔΔG mean = −0.616 kcal/mol) induce strongly stabilizing effects on the SARS-Cov-2 RBD (Table1 and Figure 2B). In all RBD mutations, G431W introduced the highest positive folding energy change at 55.323 kcal/mol, and all mutations in G431 could highly reduce the protein stabilities of RBD and full-length S (Figure 2). A397L has the smallest folding free energy change at −3.582 kcal/mol and can increase RBD stability. But many mutations in A397 (ΔΔG mean = 1.563 kcal/mol) have positive ΔΔG values and can decrease RBD stability (Figure 2B).

### Effects of RBD Mutations on RBD-ACE2 binding affinity (ΔΔΔG)

Missense mutations in RBD region could change the key interaction site and affect the binding affinity of RBD and ACE2. We calculated the binding energy changes (ΔΔΔG) of total RBD 3,705 mutations in the crystal structure of RBD-ACE2 complex. Of 3,705 mutations, 263 (7.1%) mutations can decrease (ΔΔΔG > 0.5 kcal/mol) and only 48 mutations (1.3%) can increase (ΔΔΔG < −0.5 kcal/mol) the binding affinity of RBD-ACE2 complex. Most of the mutations have limited effects (−0.5 <= ΔΔΔG <= 0.5 kcal/mol) on protein-protein interaction. As shown in Figure. 1B, 3.8% mutations have small effects on destabilizing (0.1 < ΔΔΔG <= 0.5 kcal/mol), while 3.4% mutations have small effects on stabilizing (−0.5 <= ΔΔΔG < −0.1 kcal/mol) RBD-ACE2 binding affinity. Most of the mutations (84.3%) have no effects (−0.1 <= ΔΔΔG < 0.1 kcal/mol) on RBD-ACE2 interaction.

The key residues and mutations on RBD-ACE2 binding affinity are listed in Figure 3 and Table 2. G496 has the maximum ΔΔΔG mean in RBD-ACE2 binding affinity at 4.694 kcal/mol. The mutations in this site, such as G496W and G496Y induced the first largest and second-largest binding energy change at 17.418 kcal/mol and 13.420 kcal/mol in all RBD mutations, respectively. Interestingly, G496’s neighbor residues F497 has minimum ΔΔΔG mean at −0.476 kcal/mol. Many mutations including F497W (ΔΔΔG = −0.976 kcal/mol) can reduce the RBD-ACE2 interaction. These two residues are located in a 10-residues sequence Y495**GF**QPTNGVG504. Many mutations and residues in this short motif have significant effects on RBD-ACE2 binding affinity. For example, G504 and Y495 have the second and third smallest ΔΔΔG mean values at −0.245 kcal/mol and −0.209 kcal/mol, respectively. The largest negative binding free energy change is introduced by N501E at −2.490 kcal/mol. Mutations N501L (ΔΔΔG = −2.397 kcal/mol), N501M (ΔΔΔG = −2.273 kcal/mol), N501K (ΔΔΔG = −2.134, kcal/mol) in the same position can increase the RBD-ACE2 binding affinity. In contrast, G502 has the second largest positive ΔΔΔG mean at 2.922 kcal/mol and G502P (ΔΔΔG = 11.767 kcal/mol) has the highest destabilizing effect on RBD-ACE2 complex.

**Table 2.**
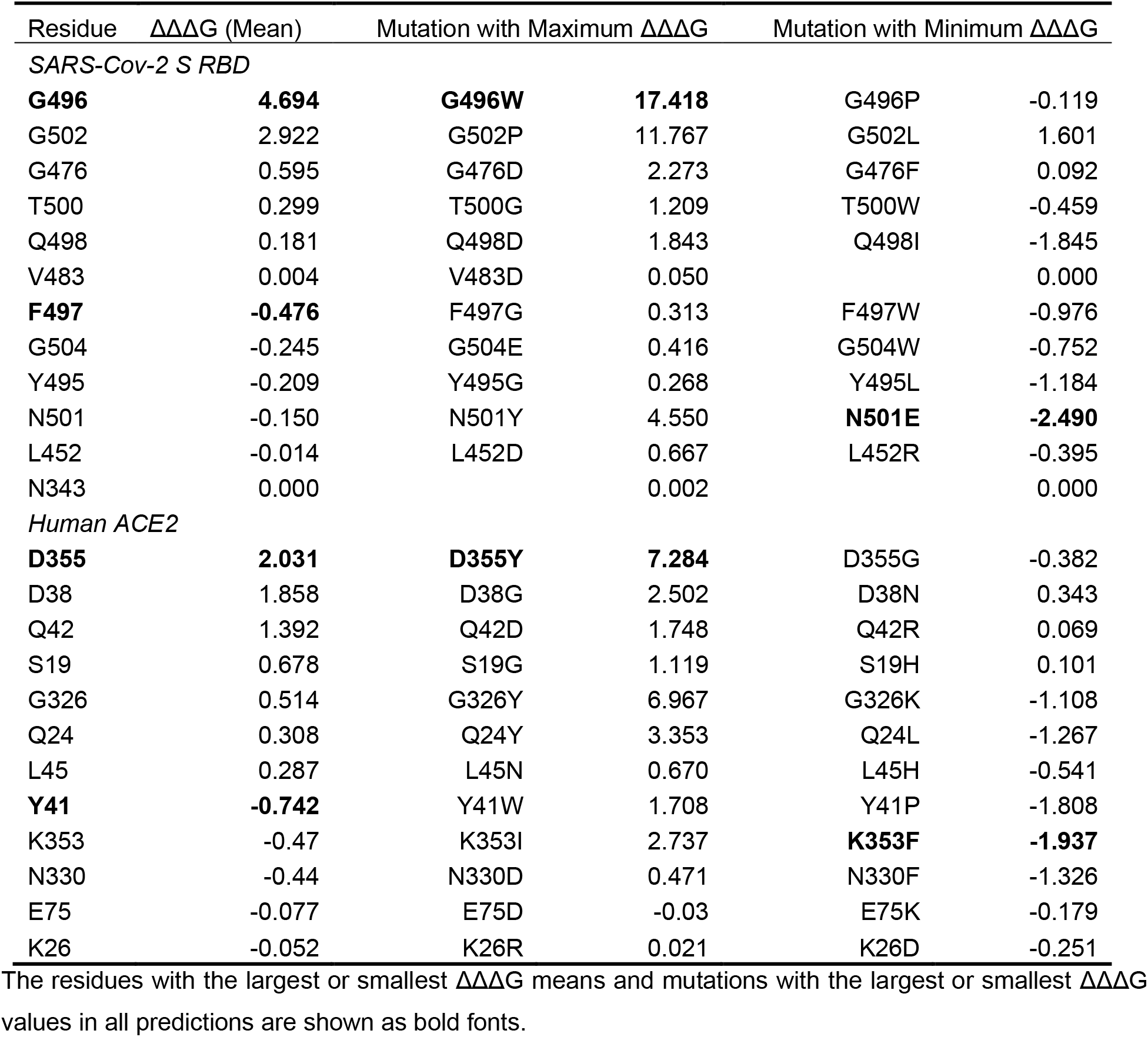
Effects of key residues and mutations on RBD-ACE2 interaction.

**Figure 3.**
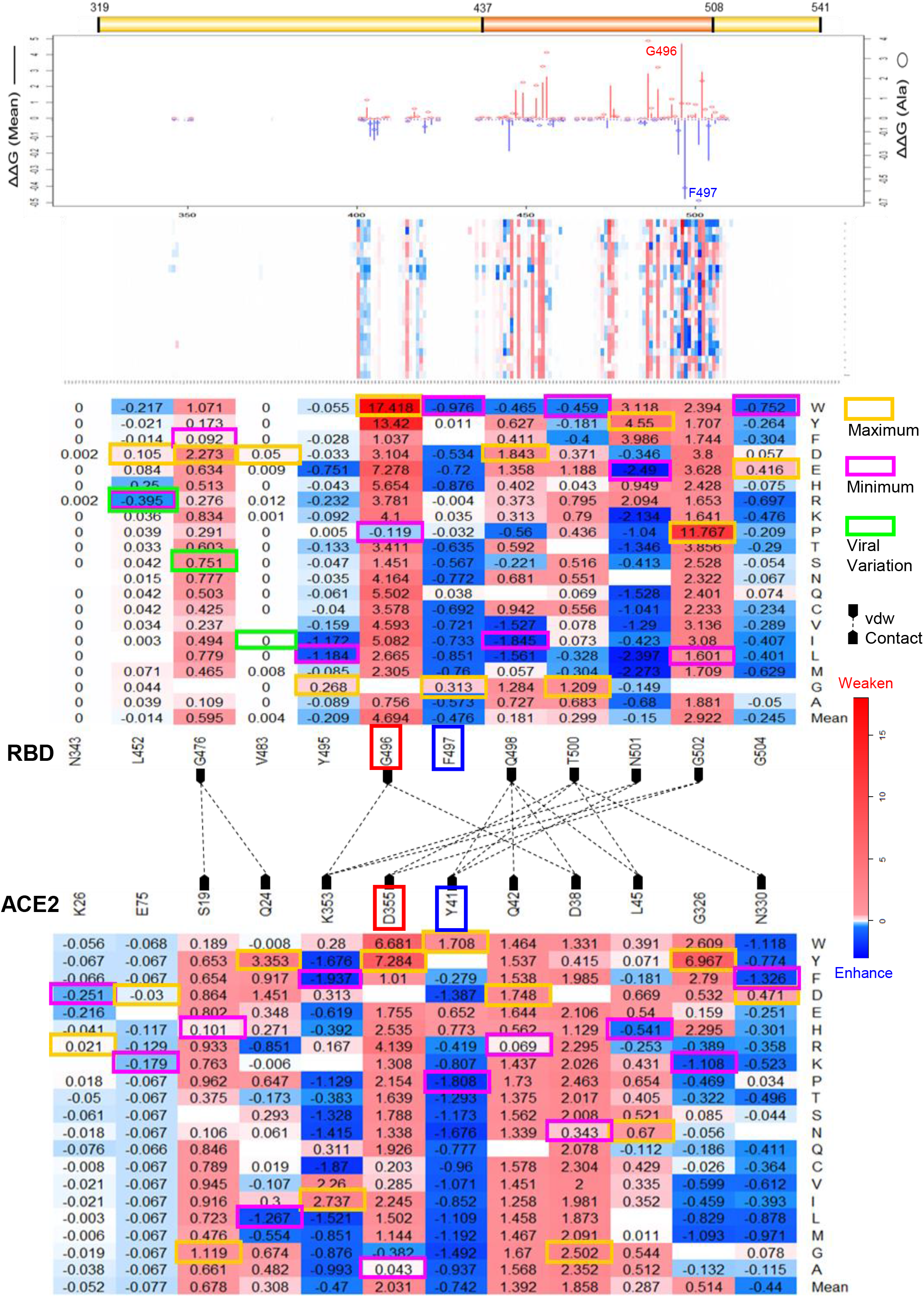
Effects of SARS-Cov-2 RBD residues and mutations on RBD-ACE2 interaction. Line chart summarizes the binding energy changes for ΔΔΔG mean of residues (bar) and ΔΔΔG of substitutions to Alanine (circle) in SARS-Cov-2 RBD residues. Heatmaps show the ΔΔΔG of all RBD mutations and the mutations in key residues. The RBD and ACE2 that have vdw contacts are marked and connected with dash lines. Maximum (yellow) and Minimum (magenta) ΔΔΔG values are labelled for each residue position. The ΔΔΔG values of viral variations are shown in green boxes. The key residues G496 and F497 are marked in the line chart and heatmaps.

### Effects of ACE2 Mutations on RBD-ACE2 Interaction

The human ACE2 mutations located in RBD-ACE2 interface could affect ACE2 interaction with SARS-Cov-2 RBD. We then scrutinized the effects of total 11,324 mutations in ACE2 chain of crystal structure of RBD-ACE2 complex (PDB ID: 6lzg). 10,729 (94.7%) ACE2 mutations have no effects (−0.1 <= ΔΔΔG < 0.1 kcal/mol) on RBD-ACE2 binding affinity (Figure. 1B). Of 11,324 mutations, 210 (1.9%) mutations can decrease (ΔΔΔG > 0.5 kcal/mol) the binding affinity and 192 (1.7%) mutations have small effects (0.1 < ΔΔΔG <= 0.5 kcal/mol) on destabilizing RBD-ACE2 complex. In contrast, only small portions, 63 mutations (0.6%) with ΔΔΔG < −0.5 kcal/mol and 130 mutations (1.1%) with −0.5 <= ΔΔΔG < −0.1 kcal/mol, can increase the binding affinity.

As shown in Table 2, the mutations at residues D355, D38 and Q42 induce strong destabilizing effects on RBD-ACE2 binding affinity. Residue D355 (ΔΔΔG mean = 2.031 kcal/mol) and mutation D355Y (ΔΔΔG = 7.284 kcal/mol) in this position have maximum effects on destabilizing the RBD-ACE2 complex. However, the mutations at residues Y41, K353 and N330 can increase the RBD-ACE2 binding affinity. Residue Y41 (ΔΔΔG mean = −0.742 kcal/mol) and mutation D355Y (ΔΔΔG = −1.808 kcal/mol) can increase the RBD-ACE2 interaction. K353F has the smallest binding free energy change at −1.937 kcal/mol, and K353C and K353Y can also increase the binding affinity of the RBD-ACE2 complex. Overall, many mutations have been identified by our studies to either enhance or attenuate the interaction between ACE2 and RBD.

### Effects of viral variations on spike protein stability and RBD-ACE2 binding affinity

We also searched and listed the viral mutations that occurred naturally in the strains of SRS-CoV-2. As of 04/30/2020, 384 experimentally identified viral missense variants have occurred in 325 positions (25.5%) of total 1273 amino acids of SARS-Cov-2 full-length S protein. Of 384 experimentally identified variants, 237 mutations can be mapped to the cro-EM structure of SARS-Cov-2 full-length S protein. We examined effects of viral missense variations on S protein stability. As shown in the Figure 1A, 32.6% viral mutations have strong effects and 11.8% mutations have moderate destabilizing effects on S protein. Remarkably, there are more viral mutations (18.1%) that have moderate stabilizing effects (−2.5 <= ΔΔG < −0.5 kcal/mol) compared to those of all computationally predicted mutations (10.4%) in the entire full-length S protein. In addition, more mutations with no effects (−0.5 <= ΔΔG < 0.5 kcal/mol) were observed in viral variations (36.3%) compared to those of all mutations (27.8%). Of 384 viral variations, 47 mutations occur in 40 positions of RBD region and 40 mutations can be mapped to the crystal structure of RBD. As shown in Figure 1B, we observed less viral mutations (2.3%) can reduce the RBD-ACE2 binding affinity (ΔΔΔG > 0.5 kcal/mol) on RBD, compare to those of all mutations (7.1%). We did not find viral mutations with ΔΔΔG < −0.5 kcal/mol. However, there are more viral mutations (9.3%) that have small effects (−0.1 < ΔΔG <= −0.5 kcal/mol) on stabilizing RBD-ACE2 complex, compared to those of all mutations (3.4%).

We investigated the effects of common viral variations and the mutations with significant effects (Table 3), The most common variation, D614G in 5703 virus strains, has stabilizing effects on the S protein (ΔΔG = −0.7838 kcal/mol). Similarly, D936Y (ΔΔG = −0.3042 kcal/mol) in 37 strains and H49Y (ΔΔG = −1.902 kcal/mol) in 25 strains can make S protein more stable. However, G1124V (ΔΔG = 3.5945 kcal/mol) in 24 strains and S943I (ΔΔG = 0.2988 kcal/mol) in 26 strains reduce the stability of S protein. Mutations of S50L (ΔΔG = −2.614 kcal/mol), T724I (ΔΔG = −2.590 kcal/mol), and T240I (ΔΔG = −2.476 kcal mol) have strong stabilizing effects on SARS-Cov-2 full-length S protein. In RBD region, the largest folding energy change takes place in S438F (ΔΔG = 18.399 kcal/mol) for S stability. Consistently, S438F has the maximum ΔΔG at 10.980 kcal/mol for RBD stability. In contrast, V341I has the minimum folding free energy change at −1.256 kcal/mol and can increase RBD stability. V483A in 24 strains have small effects (ΔΔG = −0.196 kcal/mol) and V367F in 13 strains has moderate stabilizing effects (ΔΔG = −0.597 kcal/mol) on RBD. But G476S in 8 strains, A520S in 7 strains and Q414E in 6 strains can destabilize the RBD region. We further investigated the effects of virus mutations on RBD-ACE2 binding affinity. The binding energy changes caused by common variants V483A and V367F are close to 0, meaning that these mutations have no effects on binding affinity. We observed that G476S can decrease RBD-ACE2 binding affinity (ΔΔΔG = 0.751 kcal/mol), but L452R has small stabilizing effect (ΔΔΔG = −0.395 kcal/mol) on RBD-ACE2 complex.

**Table 3.**
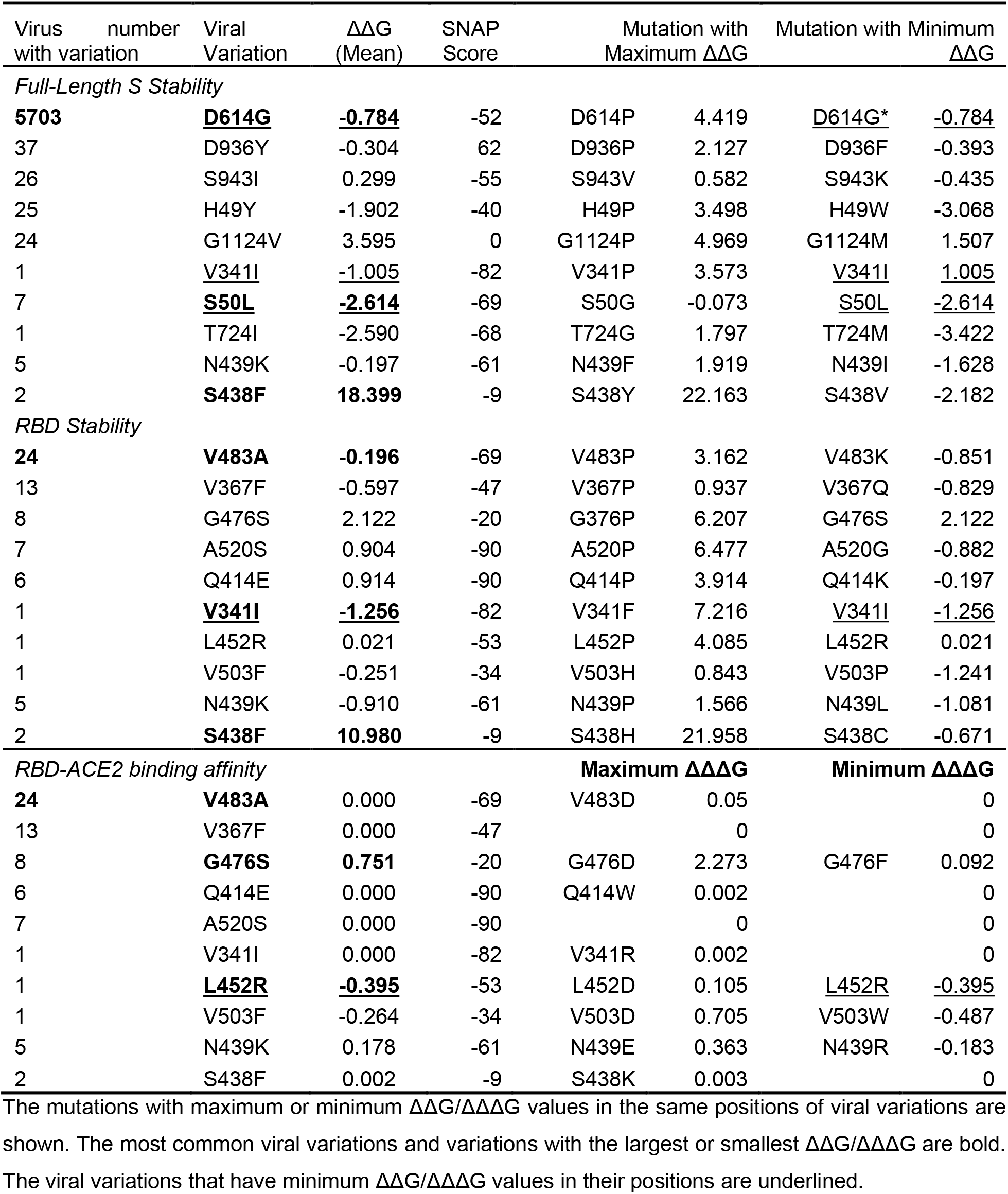
Effects of viral variations on SARS-Cov-2 S stability and RBD-ACE2 interaction.

### Mutation Pathogenicity of SARS-Cov-2 S and ACE2

We applied SNAP (26) to analyze the mutation pathogenicity of all mutations of S and ACE2 proteins based their protein sequences. We generated predictions for 15,295 mutations in human ACE2 and 24,187 mutations in SARS-Cov-2 full-length S including 4,237 mutations in its RBD region. As shown in Figure 4A, of 24,187 SARS-Cov-2 S mutations, 12, 678 (52.42%) mutations have damaging effects on S function and 11,509 (47.58%) mutations are predicted as neutral variations. In RBD region, the distribution of damaging mutations is decreased to 47.01%. Interestingly, most of viral variations (75%) have neutral effects on S function and only 25% are predicted as deleterious mutations. In human ACE2 protein, 9,628 (62.9%) mutations are predicted as deleterious variations and 5,667 (37.05%) mutations have neutral effects on protein function. The heatmap of hACE2 also suggests that most of the mutations have damaging effects on ACE2 function (Figure 4B).

**Figure 4.**
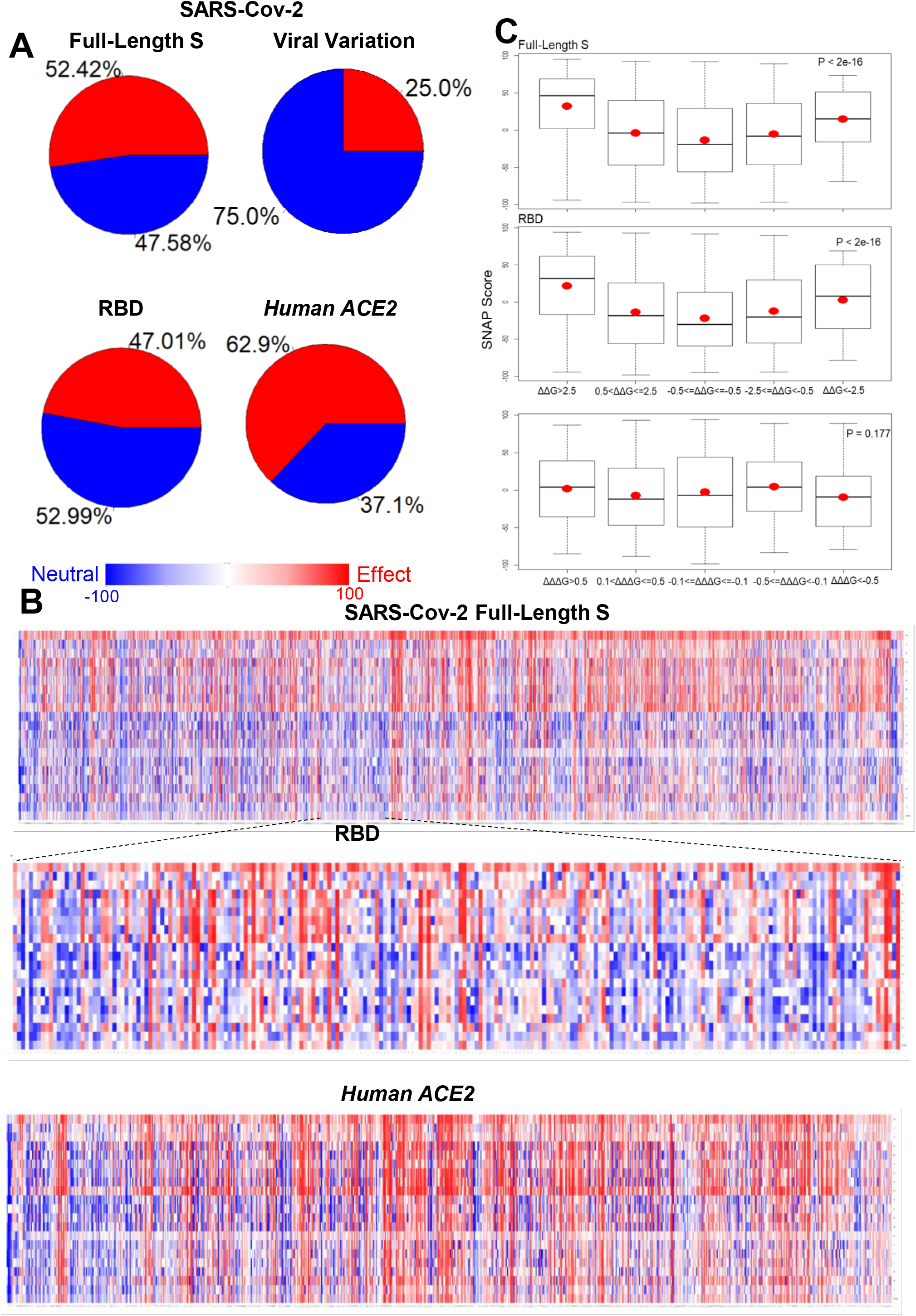
Mutation pathogenicity analysis. (**A**) Pie charts summarize the contribution of neutral (blue) and damaging (red) all mutations and viral variations in SARS-Cov-2 Full-length S and those mutations in RBD and human ACE2. (**B**) Heatmaps of mutation pathogenicity for SARS-Cov-2 Full-length S, its RBD region and human ACE2. (**C**) Boxplots for the SNAP scores of mutation groups with different folding (ΔΔG) and binding (ΔΔΔG) energy changes. The mean values are shown as red dots.

Protein destabilization is a common mechanism by which mutations cause human diseases. Studies suggested that up to 80% of the disease-causing missense mutations may lead to protein destabilization (27). The effect of a mutation on protein function can be related to its impact on protein stability changes (28). As shown in Figure 4C, we observed the SNAP scores of mutations with strong effects (ΔΔG > 2.5 or ΔΔG < −2.5 kcal/mol) on the stabilities of full-length S and RBD are higher than the mutations with moderate effects (0.5<ΔΔG <= 2.5 or −2.5 =< ΔΔG < −0.5 kcal/mol). The mutations with no effects (−0.5 <= ΔΔG <= 0.5 kcal/mol) have the lowest average SNAP scores. The differences are statistically distinguishable among these groups in SARS-Cov-2 S and RBD (Both P-value < 2e-16). Bromberg and Rost found that the correlations between mutation damaging effects and FoldX folding energy changes are strong for the mutations with significant of ΔΔG values (28). Consistent with this finding, our results suggested that the folding energy change (ΔΔG) can be used to distinguish the damaging and neutral mutations in SARS-Cov-2 S and RBD. However, we did not find the statistically difference for the mutation groups with different effects on RBD-ACE2 binding affinity (P-value = 0.177), which suggests that binding energy changes (ΔΔΔG) is not a good predictor for identifying the deleterious mutations in SARS-Cov-2 RBD.

## DISCUSSION

S protein stability is critical in producing therapeutic antibodies against current pathogenic coronaviruses (29). Pallesen et al. rationally designed a prefusion-stabilized MERS-CoV S 2P protein by introducing two consecutive proline mutations V1060P and L1061P (30). We calculated the folding energy changes of mutations in residues V1060 and L1061 of MERS-CoV wide type structure (Supplementary Figure S2). V1060P and L1061P have the minimum ΔΔG values at −2.225 kcal/mol and −0.821 kcal/mol in their positions, respectively. In addition, many mutations in V1060 (ΔΔG mean = −0.47 kcal/mol) induce highly stabilizing effects on the MERS S stability. These computational energy calculations are consistent with the findings in experimental structure analysis (30). Even though the 2P positions are not conserved among MERS-Cov, SARS-Cov and SARS-Cov-2 S amino acid sequences (Supplementary Figure S2), K986 and V987 of SARS-Cov-2 S and K968 and V969 of SARS-Cov S at equivalent positions share similar structures. All these residues are located in the linker between two helices. The folding energy calculations of K986P and V987P of SARS-Cov-2 S and K968P and V969P of SARS-Cov S show all these proline substitutions can increase the S protein stabilities. SARS-Cov-2 mutations K986P and V987P have the minimum ΔΔG values at −0.839 kcal/mol and −1.999 kcal/mol, respectively. Most mutations in V987 (ΔΔG mean = −0.334 kcal/mol) can stabilize the SARS-Cov-2 S protein. The experimental study showed that MERS-Cov 2P protein can stay in the prefusion conformation and retain high binding affinity of RBD to its receptor and various neutralizing antibodies (30). Thus, we investigated the SARS-Cov-2 residues and mutations that have significant effects on protein stability and RBD-ACE2 binding affinity.

### Residues S514 and G431 are among the most important amino acids for SARS-Cov-2 S and RBD stability

We wonder whether any other residues and mutations can induce higher stabilizing effects on SARS-Cov-2 S protein. As shown in Table 1 and Figure 2, various mutations in residues S514, S735, S50, V976 and T385 can highly increase the stability of SARS-Cov-2 full-length S, and many mutations in in S514, T385 and N394 have strong effects on RBD stability. S514 has the minimum ΔΔG mean value on both full-length S (ΔΔG mean = −1.54 kcal/mol) and RBD region (ΔΔG = −1.658 kcal/mol). The folding energy calculations to all possible mutations in this position show that S514F induce the strongest stabilizing effect on both full-length (ΔΔG = −3.304 kcal/mol) and RBD region (ΔΔG = −3.390 kcal/mol). S514 is located in a beta strand of RBD (Figure 5). Serine (S) is a polar amino acid, and phenylalanine (F) is a hydrophobic residue and aromatic amino acid. A serine to phenylalanine substitution can make this residue buried inside the protein core and increase the protein stability. We observed the significant stabilizing effects (ΔΔG < −3 kcal/mol) of the substitutions to other hydrophobic acids such as methionine (M) and leucine (L) and aromatic amino acids including tryptophan (W) and tyrosine (Y) on both full-length S and RBD.

**Figure 5.**
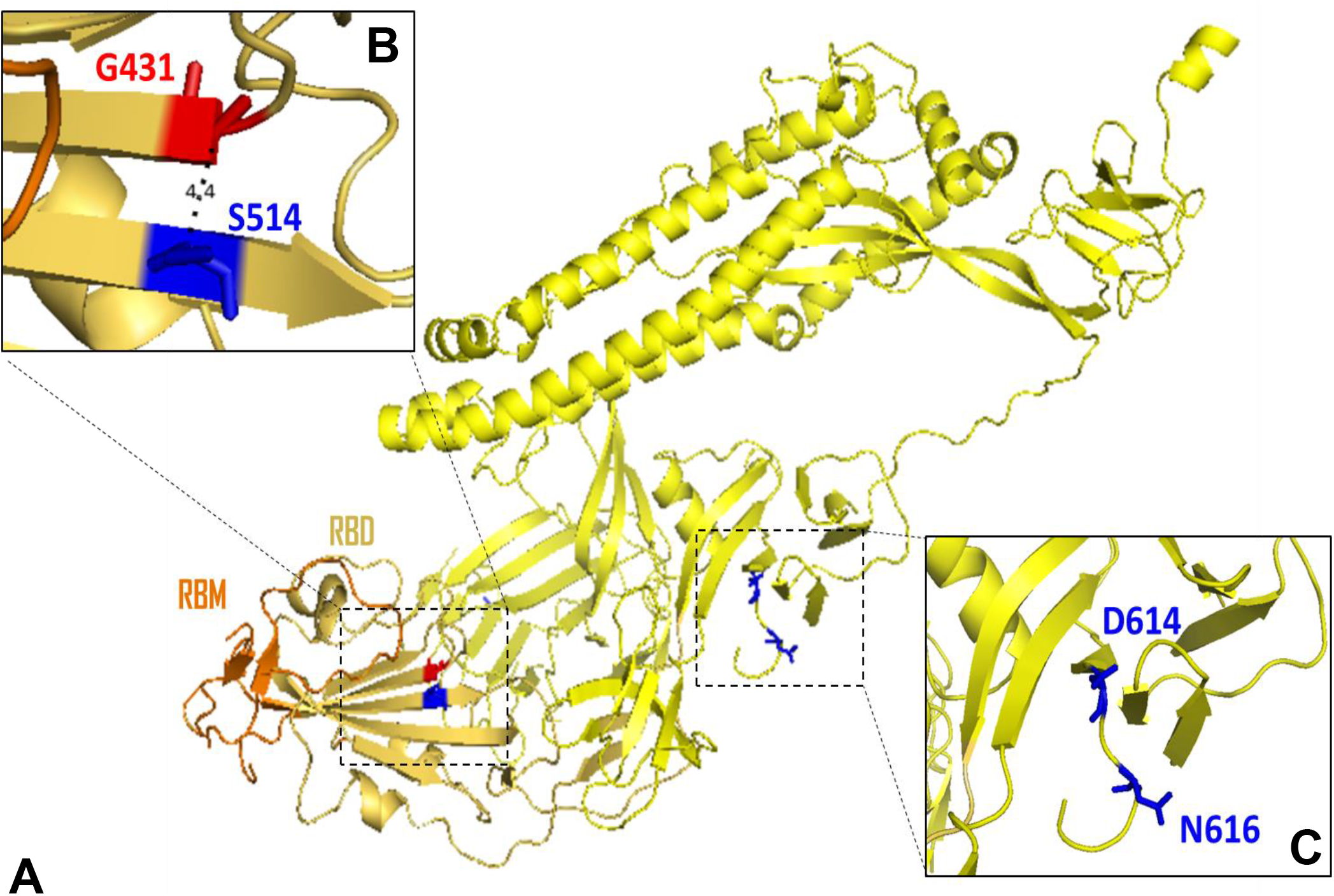
(**A**) Structural representation of key residues altering protein stability in SARS-Cov-2 full-length S (yellow). (**B**) G431 and K514 are spatial residues (4.4 Å) located in RBD (yellow orange) and closed to RBM (orange). (**C**) Viral common variation D614G located in D614 is closed to N-linked glycosylation site N616.

Destabilizing mutations can be damaging for protein function (Figure 4C) and residues with high destabilizing effects could be important active sites of spike proteins. We observed that all mutations at glycine residues G431, G648, G35 and G526 residues can destabilize both SARS-Cov-2 full-length S and RBD region. Glycine (G) is the smallest amino acid, mutations to any other large amino acids will result in unfavorable conformation changes and make protein unstable. Among all destabilizing mutations, G431W introduced the highest folding energy change on full-length S (ΔΔG = 59.92 kcal/mol) and RBD (ΔΔG = 55.32 kcal/mol), indicating this mutation can strongly reduce the protein stability. Tryptophan (W) is an aromatic amino acid. A glycine to tryptophan substitution can interact with other aromatic or positively charged residues. Interestingly, the G431 and S514 are very close structurally. The distance from alpha carbon of G431 to alpha carbon of S514 is only 4.4 Å (Figure 5B). This finding indicates the interaction between two key residues is critical for G431 and S514 SARS-Cov-2 S and RBD stability. Notably G431 is located in the upstream region of RBM and S514 is closed to the end of RBM (Figure 6A). These two residues may have an impact on RBM stability. G431 and S514 are conserved in SARS-Cov S sequence (Figure 6A). The folding energy calculations showed that all mutations in SARS-Cov residue G418, which at the equivalent position of G431 in SARS-Cov-2, can also reduce the S stability (Figure 6B). Similarly, most of the mutations in S500 of SARS-Cov have stabilizing effects like those in its conserved position S514 in SARS-Cov-2. The structure alignment of SARS-Cov-2 S and SARS-Cov S alignment suggests similar structures between two S proteins. The distance from G418 to S500 in SARS-Cov is increased to 5.3 Å, which still make interactions between two residues. These results indicate that the mechanisms of these spatial residues affecting the stabilities of S proteins are similar in SARS-Cov and SARS-Cov-2. However, our predicted mutations on G431 and S514 have not been seen in the literatures. Future experimental investigations should be focused on their topological interaction and biological functions.

**Figure 6.**
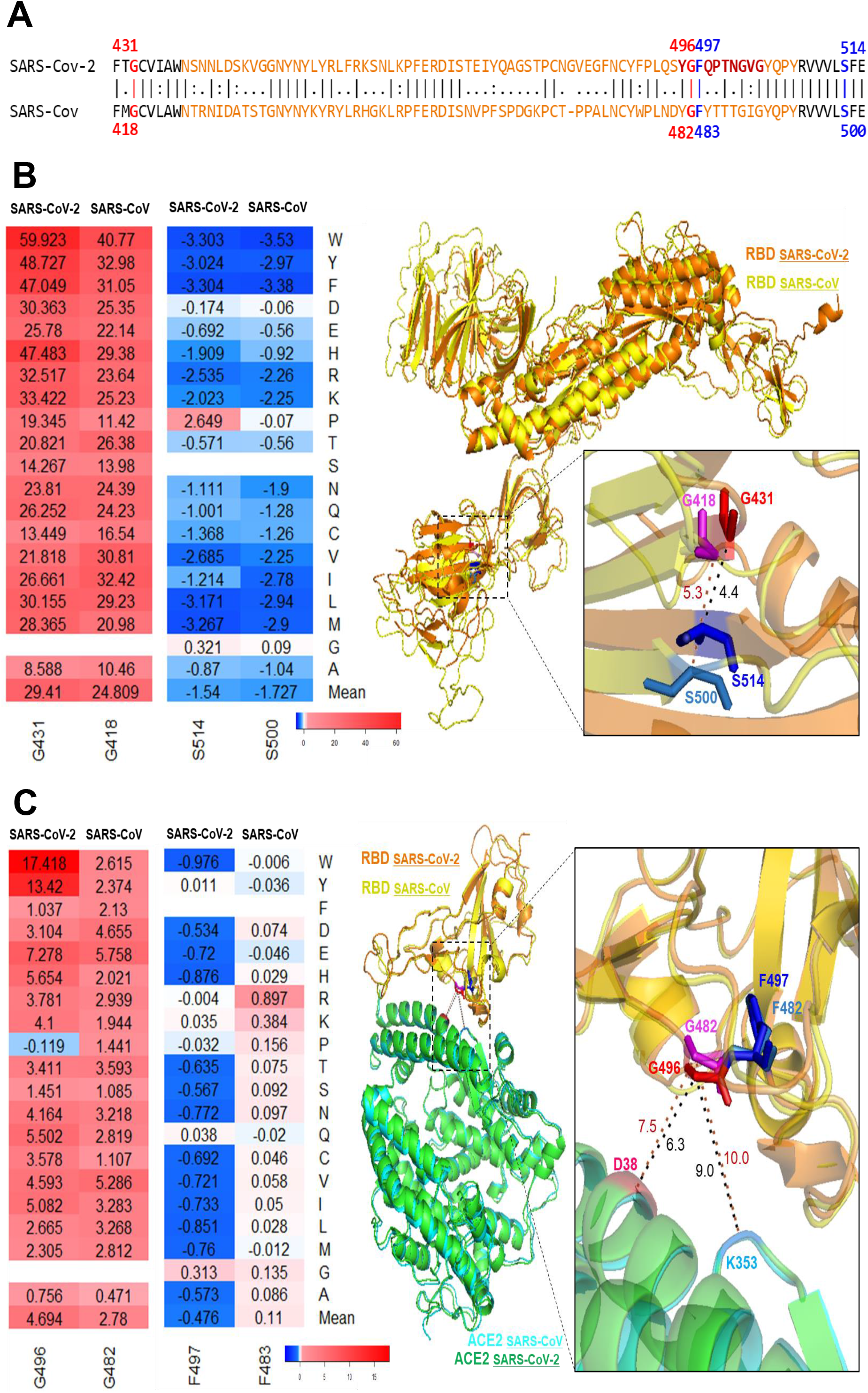
Comparison between SARS-CoV-2 and SARS-CoV. (**A**) Sequence alignment for the SARS-CoV-2 and SARS-CoV. G431 and S514 are closed to RBM (orange). G496 and F497 are located in the interacting motif Y495**GF**QPTNGVG504 (brown). These four residues are conserved in SARS-CoV. (**B**) Superimposition of the SARS-CoV-2 S (orange) and SARS-CoV S (yellow). Heatmap and structural representation of SARS-CoV-2 residues G431 (red) and S514 (blue) and SARS-CoV residues G418 (magenta) and S500 (marine). (**C**) Superimposition of the SARS-CoV-2 RBD (orange) with its ACE2 (green) and SARS-CoV RBD (yellow) with its RBD. Heatmap and structural representation of SARS-CoV-2 residues G496 (red) and F497 (blue) and SARS-CoV residues G482 (magenta) and F482 (marine). The distances to ACE2 residues D38 (hot pink) and K353 (sky blue) are measured.

### Stability effects of mutations in Glycosylation sites

Glycosylation, a post-translational modification, plays important roles in viral pathobiology and host immune responses (31). It regulates the S protein folding, increases its stability and has wide-ranging roles in viral pathobiology, including mediating, and shaping viral tropism (9). Glycosylation sites could shield specific epitopes in the spike protein from antibody recognition and facilitate immune evasion. SARS-CoV-2 S includes 22 N-linked glycosylation sequons and these oligosaccharides are critical for S protein folding (5). Watanabe et al. observed the oligomannose-type and complex-type glycans in 22 N-linked glycosylation sites using site-specific mass spectrometric (21). Of 22 N-linked glycosylation sites, 15 sites could be mapped to full-length S structure (Figure 7). N122 with oligomannose-type glycan introduces the minimum average change in protein stability at −0.452 kcal/mol. Most of the mutations in the N-linked glycosylation sites with complex-type glycans, N616 (ΔΔG mean = −0.433 kcal/mol) and N1134 (ΔΔG mean = −0.398 kcal/mol), can highly increase the S protein stability. N801D has the largest negative folding free energy change at −1.89 kcal/mol, but other mutations in this site with complex-type glycan have positive ΔΔG values. The similar mutation effect pattern is also observed in N1098 with complex-type glycan, which has the highest destabilizing effects (ΔΔG mean = 0.749 kcal/mol) among all N-linked glycosylation sites. There have been 4 observed viral mutations (N74K, N149H, N603K and N1194S) to N-linked glycosylation sites. We can calculate the folding energy change of N603K at −0.5521 kcal/mol, indicating this mutation could increase full-length S protein stability. Significantly, N603W in this position has the minimum ΔΔG at −0.92 kcal/mol. The folding energy change (ΔΔG) in this N-linked glycosylation site ranges from −0.831 kcal/mol of N343W to 1.117 kcal/mol of N343V. We also investigated two validated O-linked glycosylation sites T323 and S325. As shown in Figure 7, most of the mutations in both sites can increase the stability of SARS-CoV-2 full-length S. T323 has ΔΔG mean at −0.322 kcal/mol and T323F induce the highest stabilizing effect (ΔΔG = −1.246 kcal/mol). One viral mutation T323I was identified on this site and this mutation can increase the protein stability (ΔΔG = −0.827 kcal/mol). Remarkably, both viral mutations N603K and T323I are not the mutations introducing the strongest stabilizing effects. Of 22 N-linked glycosylation sites, only N343 with complex-type glycan can be mapped to the RBD-ACE2 complex structure. The mutations in N343 have no effects (ΔΔΔG = 0 or 0.002 kcal/mol) on binding affinity (Figure 3).

**Figure 7.**
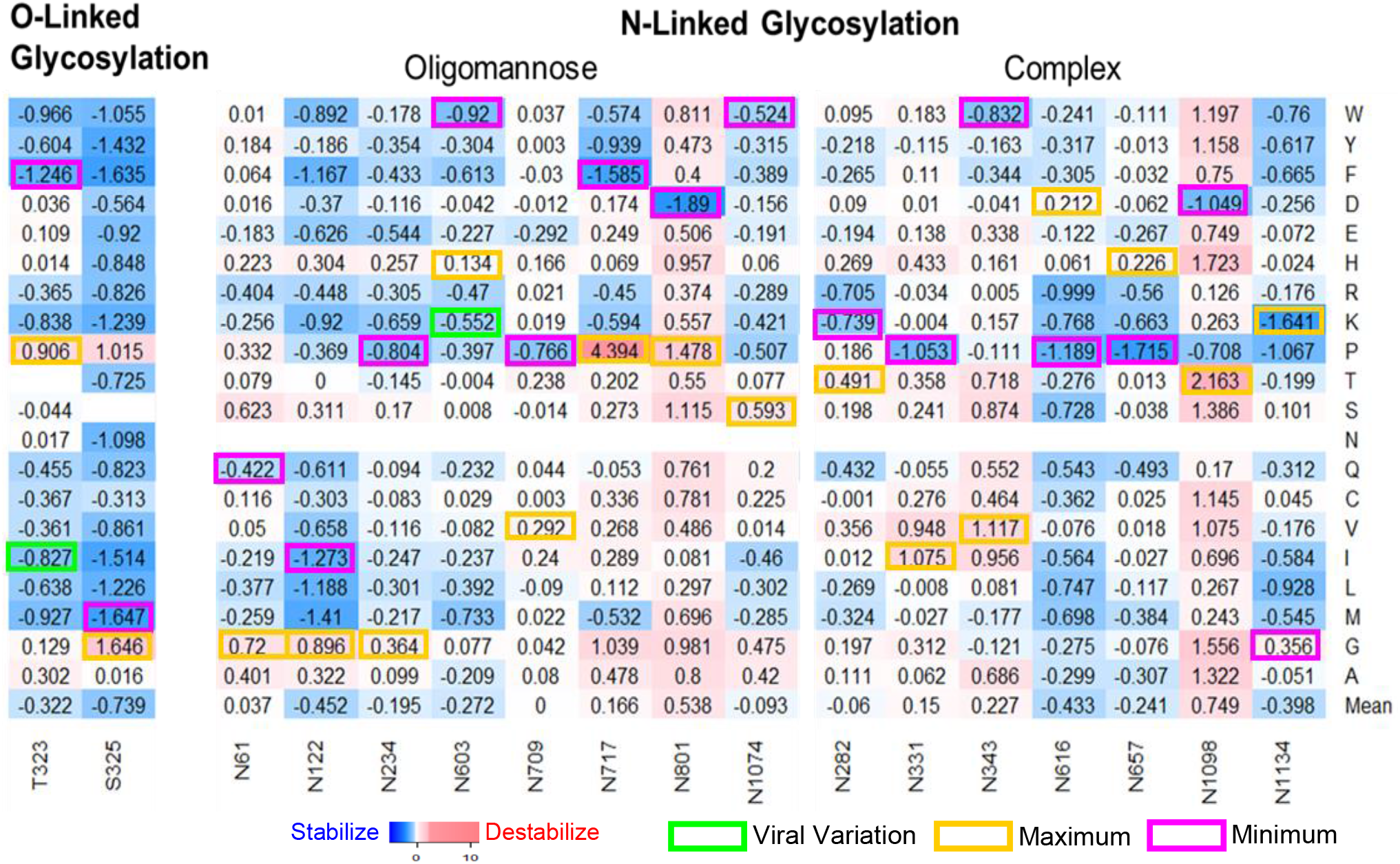
Heatmaps of mutations in glycosylation sites. Maximum (yellow) and Minimum (magenta) ΔΔΔG values are labelled for each N-linked or O-linked glycosylation site. The ΔΔΔG values of viral variations are shown in green boxes.

### Key residues in SARS-Cov-2 RBD and human ACE2 are involved in an interaction network important for RBD-ACE2 binding

Most of the mutations in SARS-Cov-2 RBD residue G496 (ΔΔΔG mean = 4.694 kcal/mol) and human ACE2 residue D355 (ΔΔΔG mean = 2.031 kcal/mol) can reduce the RBD-ACE2 interaction (Figure 3). In contrast, RBD residue F497 (ΔΔΔG mean = −0.476 kcal/mol) and ACE2 residue Y41 (ΔΔΔG mean = −0.742 kcal/mol) have the smallest ΔΔΔG mean values and many mutations in these sites can enhance RBD-ACE2 binding affinity. Moreover, the minimum binding free energy change is introduced by mutations in RBD residue N501 (N501E ΔΔΔG = −2.490 kcal/mol) and ACE2 residue K353 (K353F ΔΔΔG = −1.937 kcal/mol). We wonder whether these residues could form an interaction network (Figure 8). We investigated the binding energy changes induced by mutations in 21 RBD residues and 22 ACE2 residues that make van der Waals (vdw) contacts (32). We observed many mutations in these contact residues could alter RBD-ACE2 binding (Supplementary Figure 3). In the RBD interacting motif (Y495**GF**QPTNGVG504), RBD residue G496 with maximum ΔΔΔG mean has contacts with ACE2 residues D38 and K353 (Figure 8C). F497 with maximum ΔΔΔG mean has no vdw contacts with ACE2 residues. However, its neighbor residue Q498 has vdw contacts with residues D38, Q42, L45 as well as Y41 with minimum ΔΔΔG mean in ACE2 (Figure 8D). RBD residues T500 and N501 interact with ACE2 residues K353 and Y41 (Figure 8E), and RBD residue G502 have vdw contacts with D355 and K353 in ACE2 (Figure 8F). These viral ligand and receptor residues generate a complex network for RBD-ACE2 interaction (Figure 3 and Figure 8).

**Figure 8.**
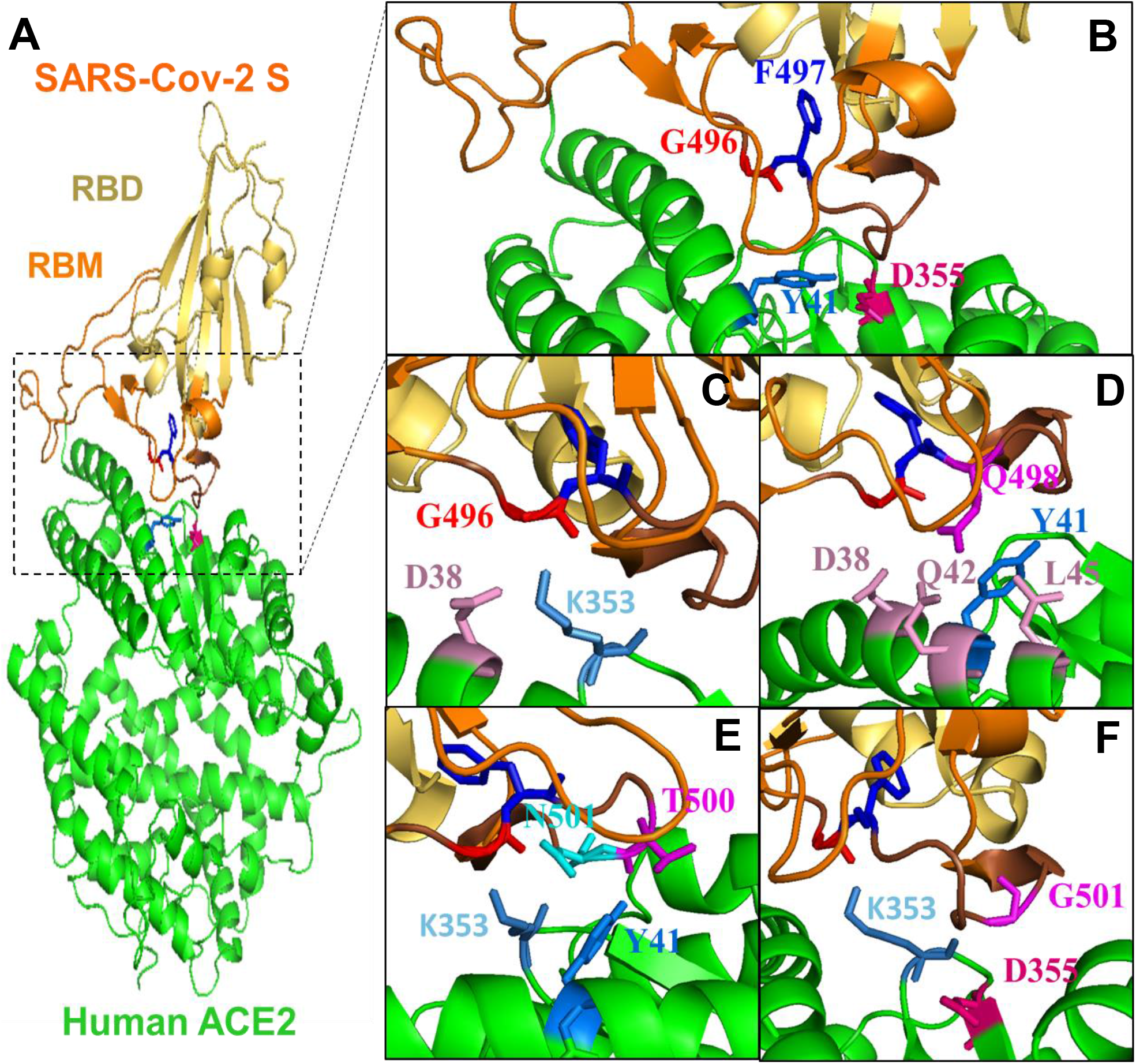
(**A**) Structural representation of key residues altering the binding affinity between SARS-Cov-2 S (RBD: yellow orange; RBM: orange) and human ACE2 yellow. (**B**) G496 (red) and F497 (blue) are located in the interacting motif Y495**GF**QPTNGVG504 (brown) and form an interaction network with ACE2 residues D355 (hot pink) and Y41 (marine). (**C**) RBD residue G496 with has vdw contacts with ACE2 residues D38 (pink) and K353 (sky blue). (**D**) Q498 has contacts with residues D38, Q42, L45 (pink) as well as Y41 (marine). (**E**) RBD residues T500 (magenta) and N501 (cyan) interact with ACE2 residues K353 (sky blue) and Y41(marine). (**F**) RBD residue G502 (magenta) have vdw contacts with D355 (hot pink) and K353 (sky blue) in ACE2.

SARS-Cov-2 S neighbor residues G496 and F497 introduce the most significant effects on RBD-ACE2 binding. Of 19 mutations in residue G496 (ΔΔΔG mean = 4.694 kcal/mol), 15 mutations can highly reduce the RBD-ACE2 binding affinity (ΔΔΔG > 2.5 kcal/mol), 3 mutations have the moderate destabilizing effects (2.5 => ΔΔΔG > 0.5 kcal/mol) and only 1 mutation G496P has small stabilizing effect (ΔΔΔG = −0.119 kcal/mol) on RBD-ACE2 complex (Figure 3). The substitutions from small glycine to other large amino acids will lead to an unfavorable change in protein interface and reduce the binding of complex. In G496’s contact residue K353 of ACE2, mutation K353F has the minimum ΔΔΔG at −1.937 kcal/mol. Most mutation in RBD residue F497 can increase the RBD-ACE binding, and mutation F497W has the minimum ΔΔΔG at −0.978 kcal/mol in this position. Both phenylalanine (F) and tryptophan (W) are aromatic amino acids which include delocalized π electrons that interact with other aromatic residues as well as with positively charged residues such as lysine (K).

SARS-Cov RBD residues G482 and F483 are in the equivalent positions of G496 and F497 of SARS-Cov-2 in the sequence and structure alignments (Figure 6A and 6C). All mutations in SARS-Cov RBD residue G482 can reduce the RBD-ACE2 binding affinity but the ΔΔΔG values are relatively small compared with those changes in G496 of SARS-Cov-2. This may be explained by that the distances from SARS-Cov residue G482 to ACE2 residues D38 and K353 are increased to 7.5 Å and 10.0 Å, compared to the distances from SARS-Cov-2 residue G496 to ACE2 residues D38 and K353 are 6.3 Å and 9.0 Å (Figure 6C). Interestingly, even though the F483 of SARS-Cov can be superimposed to F497 of SARS-Cov-2, F483 shows the different effects on RBD-ACE2 interaction. Most of the mutations in F483 of SARS-Cov have not effects on binding affinity and F483R can weaken the interaction. However, most of mutations in F497 of SARS-Cov-2 can enhance (ΔΔΔG < −0.5 kcal/mol) the RBD-ACE2 binding affinity. Previous studies suggest that SARS-Cov-2 S protein has a higher RBD-ACE2 binding affinity compared to those of SARS-CoV S (5, 10, 14). F497 may play important roles for enhancing the RBD-ACE2 interaction for SARS-Cov-2. Our binding energy calculations suggest that these residues are potential binding sites for spike protein and its receptor, any alterations in these sites may significantly change the binding affinity of RBD-ACE2 complex.

### Common viral variations tend to increase spike protein stability

We found that the most common variants D614G in 5703 strains can induce the stabilizing effects on SARS-Cov-2 full-length S at −0.7838 kcal/mol. Particularly, D614G becomes the dominant pandemic form worldwide (33). D614 form is in the Wuhan reference strain and a G-to-A base substitution result in the G614 form. The D614 mutation was first found in the strains in Germany. Korber et al. found that G614 form started its expansion in Europe and then spread quickly to many other countries in the world (33). They showed that the patients with D614G strain had higher viral loads and suggested this mutation is important for RBD binding and enhance viral infection and production. Based on our calculations of folding energy changes, D614G has the minimum ΔΔG at −0.7838 kcal/mol among all 19 possible mutations in this position (Figure 2). This suggests that G614 is the most stable form compared to other possible mutations. In addition, D614G is predicted to have neutral effect (SNAP score = −52) on protein function, which would benefit the viral S protein function (Table 3). Interestingly, we also found D614 is very closed to N-linked glycosylation site N616 (Figure 5C). Many mutations in N616 can also induce the stabilizing effects (Figure 7). Thus, D614G may enhance the fitness of SARS-Cov-2 through increasing S protein stability and participating in the N-linked glycosylation. In addition, common viral mutations D936Y in 37 strains, V483A in 24 strains and V367F in 24 strains can also induce the stabilizing effects on SARS-Cov-2 full-length S or RBD (Table 3). S50L in 7 strains and V341L have the minimum ΔΔG among all other possible mutations in their positions (Supplementary Figure 4). Moreover, we observed the percentage of mutations with stabilizing effects is higher in viral mutations (18.9%), compared to the one in all computationally predicted mutations (10.6%) (Figure 1). These results indicate that the mutations with stabilizing effects can make S protein remain sufficiently stable for its function and enhance the resistance of SARS-CoV-2. We noticed that most of the viral mutations are not substitutions with minimum and maximum folding or binding energy changes (Supplementary Figure 4). As shown in Supplementary Figure 5, the |ΔΔG| mean of residues with viral mutations is lower than those without viral mutations (P < 2.2e-16), and |ΔΔG| of viral mutations is also smaller than other computational predicted mutations (P = 0.00016) in the same positions. The mutations with significant stabilizing or destabilizing effects may interfere with the S protein function (Figure 4C). The selection pressure makes the viral mutations occurred in the residues with small effects on protein stability and maintain SARS-Cov-2 S at its normal functions for transmission.

## CONCLUSION

The emergence of the pathogenic betacoronavirus SARS-CoV-2 poses a serious threat to global health. SARS-CoV-2 S protein and its RBD and human ACE2 are critical for viral success to infection. In this study, we applied the computational saturation mutagenesis to mutate all residues in the S and ACE2 proteins to all other 19 amino acid types. We used structure-based energy calculations and sequence-based bioinformatics tools to quantify the systemic effects of missense mutations on the protein structure and function. A total of 18,354 mutations in SARS-CoV-2 spike protein were analyzed and we discovered that the majority of these mutations could destabilize the entire S protein including its RBD region. Remarkably, the amino acid substitutions in SARS-CoV-2 RBD residue G431 can decrease the spike protein stability, but the mutations in its spatial residue S514 can make the spike and RBD more stable. We analyzed 384 experimentally verified spike missense variations and identified that D614G in 5,703 virus strains can stabilize SARS-Cov-2 entire spike protein. Moreover, we showed that many mutations in N-linked glycosylation sites can increase the stability of the spike protein. In addition, we investigated 3,705 mutations in SARS-CoV-2 RBD and 11,324 mutations in human ACE2 and found that the mutations located in the interface of RBD-ACE2 complex can alter its binding affinity. Interestingly, SARS-CoV-2 neighbor residues G496 and F497 have different effects on RBD-ACE2 binding and ACE2 contact residues D355 and Y41 are critical for this protein-protein interaction. Overall, the analysis is critical for understanding the roles of missense mutations in SARS-CoV-2 S and human ACE2 proteins on the viral pathogenesis of COVID-19.

## ACKNOWLEDGEMENT

We acknowledge the Howard University Junior Faculty Writing & Creative Works Summer Academy.

## FUNDING

This work was supported by the Howard University startup funds U100193 and National Science Foundation HDR DSC Award #1924092 (ST). This project was supported (in part) by the National Institute on Minority Health and Health Disparities of the National Institutes of Health under Award Number 2U54MD007597. The project was partially supported by NIH/NIAID 1SC1AI112785-05 and R01DE028583-01 (QT). The content is solely the responsibility of the authors and does not necessarily represent the official views of the National Institutes of Health.

## AUTHOR CONTRIBUTIONS

ST and QT are PIs on the grant funding and supervised the study. ST designed the study and performed the bioinformatics analysis. ST, AS, RR, DL and QT analyzed data. ST wrote the manuscript, and all authors critically revised the manuscript.

## SUPPLEMENTARY FIGURES LEGENDS

**Supplementary Figure 1.** Folding energy changes for ΔΔG of substitutions to Alanine (left) and ΔΔG mean of residues (right) in (**A**) the cryo-EM structures of SARS-CoV-2 S in closed state and open state and (**B**) the crystal structure of RBD and cryo-EM structure of SARS-Cov-2 S RBD.

**Supplementary Figure 2.** Comparisons of 2P mutations among MERS-Cov, SARS-Cov-2 and SARS-Cov.

**Supplementary Figure 3.** Heatmaps show the ΔΔΔG of the mutations in RBD residues and ACE2 residues that make van der Waals contacts. Maximum (yellow) and Minimum (magenta) ΔΔΔG values are labelled for each residue position.

**Supplementary Figure 4.** Heatmaps of viral mutations (green) for (**A**) Full-length S stability (ΔΔG), (**B**) RBD stability ΔΔG and (**C**) RBD-ACE2 binding affinity (ΔΔΔG). Maximum (yellow) and Minimum (magenta) ΔΔG /ΔΔΔG values are labelled for each residue position.

**Supplementary Figure 5.** Boxplots for the |ΔΔG| of mean of residues with and without viral mutations (upper) and |ΔΔG| of viral mutations and other computationally predicted mutations in the same positions (lower). The mean values are shown as red dots.

## Notes

### Competing Interest Statement

The authors have declared no competing interest.

